# Diversity and prevalence of *Anaerostipes* in the human gut microbiota

**DOI:** 10.1101/2025.05.01.651700

**Authors:** Disha Bhattacharjee, Lindsey C. Millman, Meagan L. Seesengood, Anna M. Seekatz

## Abstract

Members of the class Clostridia, a polyphyletic group of Gram-positive, spore-forming anaerobes in the Bacillota (Firmicutes) phylum, are prevalent and commonly associated with beneficial functions, such as providing colonization resistance against pathogens. Despite a demonstrated value in maintaining Clostridial populations in the gut, the functional and strain diversity of most commensal Clostridial species remains understudied. Here, we isolated and characterized Clostridial isolates, focusing on the genomic diversity of *Anaerostipes*, a prevalent butyrate-producing genus within the gut microbiota. We conducted a genomic comparison across 21 *Anaerostipes* strains isolated from healthy human fecal samples (n = 5) and publicly available genomes (n = 105). Whole genome comparisons across the *Anaerostipes* genus demonstrated 12 species bins, clustering into three major functionally distinct clusters correlating with host origin. One cluster (representing mostly *A. caccae* genomes) was distinguished by possessing a complete vitamin B12 biosynthesis pathway. Variability in carbohydrate and amino acid metabolism was demonstrated within dominant species of the human microbiota (*A. hadrus, A. caccae*, and *A. hominis*). Collectively, these data indicate metabolic variance across *Anaerostipes* species that may influence coexistence within the gut environment and variably influence health.

**IMPORTANCE:** *Anaerostipes* is a genus containing species known to produce butyrate upon fermenting lactate. Despite their association with health across studies using 16S rRNA gene-based analyses, little is known about genomic variability within and across species. Our study represents one of the first analyses to define strain variability across the *Anaerostipes* genus, identifying three functional clusters and close phylogenetic distance within species found in the human gut. Major variability within species prevalent in the human gut included variable carbohydrate and amino acid metabolism genes, suggesting the ability to coexist in the gut environment.

## INTRODUCTION

The gut microbiota plays a vital role in human health by providing essential physiological and developmental functions, including training host immunity, supporting digestion of dietary intake, and regulation of neurological signaling^1^. Disruption of this ecosystem can lead to colonization of pathogenic bacteria or loss of species that support these functions, influencing various conditions or diseases that harm the host^2^. Understanding the tenets that uphold the structure of a healthy or beneficial microbiota, both at the community and individual species level, can thus guide preventative and interventional methods to modulate the gut microbiota, which may ultimately be used to treat disease.

In addition to considering how differences in species composition influence gut microbiota functions, genotypic variation across individual strains may influence microbial functional outcomes^3^. Much of our initial definitions of a healthy microbiota has been based on 16S rRNA gene-based analyses, which cannot typically distinguish taxonomic groups beyond the genus level^4^. While useful for initial surveys of microbiota, this method does cannot account for gene content or genomic differences within a given species, which can disparately influence the overall output of a community or differentially impact the host^5,6,7^. While genomic variation is well-established when considering virulence of pathogens^8^, its importance has been less considered within the context of the human gut microbiota.

Lack of information on phenotypic and genomic variation is especially true for many members of the polyphyletic class, Clostridia^9^. Clostridia encompass a broad group of spore-forming, saprophytic anaerobes. Many species within this group, such as *Faecalibacterium prausnitzii, Clostridium scindens, Coprococcus comes, Clostridium sporogenes* and others, are also frequently correlated with health^10^. Importantly, this group encompasses many species that produce metabolites of interest to human health. Lachnospiraceae such as *Coprococcus*, *Roseburia*, *Anaerostipes*, *Blautia,* and others, are known to produce short chain fatty acids (SCFAs) upon fermentation of dietary fiber and resistant starches^11,12,13,14^. The SCFA, butyrate, is a main energy source for colonocytes, and has been demonstrated to play a regulatory role in epithelial defense barrier, intestinal motility, and inflammation, making microbes producing butyrate promising candidates for future probiotics^15,16^. Clostridia also include species such as *Clostridium scindens* and *Clostridium hiranonis* that modulate bile acids initially produced by the host, which influence host physiology in diarrhea-predominant irritable bowel syndrome (IBS-D) and colonization resistance against *Clostridioides difficile* infections^17,18^. Clostridia can additionally influence host health through protein fermentation. For example, *F. prausnitzi* one of the most abundant butyrate producers, can influence gut physiology in gnotobiotic mice and restore serotonin in IBS murine models^19^. Similarly, another gut symbiont, *C. sporogenes* can produce indole propionic acid from dietary tryptophan, which can fortify the intestinal barrier^20^.

A major producer of butyrate includes the genus *Anaerostipes*^21^, a member of the Lachnospiraceae family belonging to phylum Bacillota (previously Firmicutes^22^). Initially classified solely as *Anaerostipes caccae*, the *Anaerostipes* genus now contains the major species *A. caccae, A. hadrus, A. rhamnosivorans*, and *A. butyraticus*^23,24,25,26^. Lower abundances of *Anaerostipes* have been associated with gut diseases, such as colorectal cancer^27^, inflammatory bowel disease (IBD), and irritable bowel syndrome (IBS)^28,29^. *A. hadrus* has been studied *in vitro* for its ability to produce butyrate via butyryl-CoA: acetate CoA-transferase^30^ and butyrate kinase^31^ pathways, which are also present in *A. caccae*. Most recently, *A. caccae* administration in mice was shown to be effective in reducing severe responses to allergen challenge, demonstrating a role in immune modulation and potential therapeutic response^32^. However, both genomic variability and characterization of this species’ niche within the microbiota in directing potential beneficial functions, remains less characterized.

Here, we investigate the diversity and metabolic potential of *Anaerostipes* using comparative genomics. An isolation pipeline targeting Lachnospiraceae species from the human gut consistently demonstrated the presence of *A. hadrus* and *A. caccae*. Genomic comparison to high-quality genomes from publicly available databases demonstrated host-specific variation of *Anaerostipes* species, clustering into three distinct functional clusters. In addition to exhibiting widespread butyrate-producing pathways, *Anaerostipes* also exhibited genes associated with bile acid modification, vitamin B12 production, and tocopherol cycling, expanding the potential functional contributions of *Anaerostipes* in the human gut.

## MATERIALS AND METHODS

### Isolation of Commensal Clostridia from Healthy Human Fecal Samples

This study was approved by Clemson University’s Institutional Review Board. Healthy donors were over 18, had not taken antibiotics or been diagnosed with any infections within 6 months, and were not immunocompromised or diagnosed with chronic gastrointestinal conditions. Upon receipt, fecal samples were placed under anaerobic conditions (Coy Laboratory Products, Grass Lake, MI, USA; 85% nitrogen, 10% hydrogen, and 5% carbon dioxide) and both a direct fecal streak and a 1:10 dilution fecal slurry were streaked out onto Brain Heart Infusion (BHI), BHI with the addition of fetal bovine serum (BHI+FBS) (samples CM01-CM03), BHI with the addition of rumen fluid (BHI+R) (samples CM06-CM11), taurocholate cycloserine-cefoxitin-fructose (TCCFA), reinforced Clostridia medium (RCM), and yeast casitone fatty acids (YCFA) agar plates as performed previously^33^. The fecal slurry was also used to inoculate 5 mL of each of the respective broth medias. Both the plates and broth cultures were incubated at 37°C for 24 hours. After incubation, broth cultures from each media were diluted up to 10^−6^ colonies per mL and streaked out onto their respective plate types. Unique colonies based on morphology were picked from all incubated plates over the course of several days, given a unique number, and re-streaked to purity. Once isolates were found to be pure, a single colony was used to inoculate 5 mL of the same media as the plate and grown 24-48 hours until visible turbidity was observed. Monoculture broths were divided into aliquots for initial taxonomic identification via Sanger sequencing and storage in 20% glycerol at -80 °C, preserving remaining broth for additional DNA extractions if necessary.

### DNA Extraction and Identification of Fecal Isolates

Broth aliquots were heat-extracted at 95°C for 20 minutes and prepared for polymerase chain reaction (PCR) using GoTaq (Promega; catalog no. M7132), using the 8F (5’-AGA GTT TGA TCC TGG CTC AG -3’) and 1492R (5-’ GGT TAC CTT GTT ACG ACT T-3’) primers to amplify the whole 16S rRNA gene. PCR products were cleaned up using Exo SAP-IT (Applied Biosystems, 78-200-200uL) and sent to Eton Biosciences (www.etonbio.com) for Sanger sequencing. Sequences were taxonomically identified using the NCBI, EzBioCloud, and RDP databases. Visualizations were done in R.

For 21 *A. hadrus* and *A. caccae* strains, DNA was extracted from 1.8 mL of overnight culture using the Qiagen DNeasy UltraClean microbial kit (Qiagen; catalog no. 12224-250). Extracted DNA was diluted to 10 ng/μL concentration (Qubit, Life Technologies; catalog no. Q33230) and sent to SeqCenter, Pittsburgh (www.seqcenter.com) for Illumina sequencing using the NextSeq2000 platform.

### Assessment of Lachnospiraceae in human 16S rRNA gene-based surveys

Multiple FASTA sequences of full-length 16S rRNA sequences from different Lachnospiraceae genomes that were sequenced as a part of the isolation pipeline were formatted for alignment in mothur^34^ (v1.48.0) and aligned using the SILVA database^35^ (v138.2). Previously published pairs of sequences from fecal microbiota samples representing healthy adult individuals (Supplementary Table 2) were processed in mothur using the Schloss lab standard operating procedure (SOP), aligning to the SILVA database and then classifying to the custom classifier using the classify.seqs command in mothur (cutoff = 80) as previously published^33^. The log_10_ relative abundance percentage and prevalence of various Lachnospiraceae was plotted in R for visualization.

### Whole genome assembly and phylogeny

We used a previously published workflow for assembly of all genomes listed in Supplementary Table 1^33^, also available on (https://github.com/SeekatzLab/Anaerostipes-genomes). Briefly, raw reads were quality-checked and adapter-trimmed using Trim-galore (v0.6.5)^36^, assembled using SPAdes^37^ (v3.15.5). Quast (v5.0.2) with MultiQC (v1.27.1) was used to calculate assembly statistics (Supplementary Table 1)^38,39^. Average coverage was calculated using Bowtie2 and SAMtools^40,41^. Prokka (v1.14.5) was used to annotate assemblies^42^. To verify the assembly identity, 16S rRNA gene were run through NCBI Blast and EzBioCloud. Assemblies were also mapped on to the Genome Taxonomy Database (GTDB)^43^ through GTDB-tk (v2.4.0) using classify-wf^44^. Maximum likelihood trees from the *Anaerostipes* core genome SNP sites (using SNP-sites v2.5.1) was determined by Roary and trees were created using RaXML (v8.2.12) bootstrapping 500 times^45,46,47^. AAI calculations and resultant Unweighted Pair Group Method with Arithmetic Mean (UPGMA) hierarchical clustering tree were obtained using EzAAI^48^. Trees were visualized using ggtree, ggtreeExtra and treeio packages in R^49,50,51^. Final 126 *Anaerostipes* genomes for all analysis were selected based on: N50 >25000, Total length between 2.5 Mbp and 3.7 Mbp, number of contigs < 251, checkM^52^ contamination < 5% and completeness > 95%.

### Pangenome analysis, functional enrichment, average nucleotide identity, and dereplication

Contigs from SPAdes were reformatted and annotated with the COG and KEGG database using Anvi’o ^53^ (v8.0). Anvi’o was also used to create and visualize the pangenomes, determine average nucleotide identity (ANI), and dereplicating strains within the dataset. Heap’s law was calculated in R (formulated as n = κN^γ^, where n is the pan-genome size, N is the number of genomes used, and κ and γ are the fitting parameters) and the α parameter from Power law model was calculated using micropan^54,55^. Average nucleotide identity (ANI) was computed using the anvi-compute-genome-similarity with pyANI^56,57^. Dereplication between strains was computed using anvi-dereplicate-genomes at 90, 95, 98, 99, 99.9 and 100 % similarity threshold. Functional enrichment was calculated using anvi-compute-functional-enrichment-across-genomes along with corresponding statistics^58^ and microbial metabolism is calculated using anvi-estimate-metabolism^59^. All visualization was performed using dplyr, ggplot2, and readxl packages^60,61,62^.

### SCFA, BA, toxin, sporulation, and germination genes and respective trees

Databases for *eutD*, *tdcD*, *abfD*, *ato*, *buk*, *but*, *cro*, *gcd*, *kal*, *epi/mce*, *lcdA*, *mmdA*, *mut*, *pduC*, *pduP*, *ygfH*, *bsh*, *gerAB*, *spo0A*, and *zona* were created through UniProt^63^ using the name of the gene, only bacterial taxonomy and respective appropriate length of the protein. DIAMOND v2.0.14 was used to create a Blast appropriate database and Blast (--id 70 --query-cover 80 --top 1) against the proteins from each genome^64^. Visualization of the resultant hits was done in R. For the trees for *gerAB*, *bsh*, *but*, and *buk*, corresponding hits and their associated protein sequences were aligned using the multiple sequence aligner ClustalO v1.2.4^65^. Maximum likelihood trees of the resultant alignment were generated using RAxML and visualized using ggtree in R as stated above.

### CAZyme and putative virulence

CAZymes were predicted using DBCAN (v4.1.4)^66^, using the Fasta nucleotide sequences generated from Prokka for each of the strains and visualized through R. Prokka generated nucleotide fasta files (.fna) were processed through PathoFact (v1.0)^67^ to predict virulence factors, toxins, and antimicrobial peptides and visualized through R.

### Sporulation Assay

Two strains of *A. hadrus* (CM02-05 and CM03-84) and two strains of *A. caccae* (CM03-34 and CM06-64) were streaked from frozen glycerol stocks onto BHI agar plates and incubated for 48 hours at 37°C under anaerobic conditions. Isolated colonies were picked and used to inoculate 5 mL of Clospore broth media^68^. These were then incubated at 37°C for 14 days to induce sporulation. After this time, each strain was observed under phase contrast microscopy to confirm the presence or absence of endospores. Each sample was observed under 1000X oil immersion on a Leica phase contrast microscope (Leica DM750 Phase Contrast), and pictures were taken with a Leica ICC50W camera, and the Leica Acquire software.

### Data Availability

All analyses were done through the Clemson University’s high performance computing center (HPCC), Palmetto^69^ and RStudio. All raw sequence data and associated information has been deposited in the NCBI Sequence Read Archive under BioProject PRJNA1241798. All code used to analyze data are available at https://github.com/SeekatzLab/Anaerostipes-genomes.

## RESULTS

### *Anaerostipes* are prevalent members of the human gut microbiota

We screened nine fresh human fecal samples as part of a larger project aimed at the cultivation of various gut commensal bacteria (Figure 1). Sanger sequencing of the full 16S rRNA gene from morphologically distinct colonies revealed a total of 1,038 isolates, representing 265 unique species. Of these, five phyla were identified, with representatives of the Firmicutes (88.05%), Proteobacteria (3.95%), Actinobacteria (3.85%), Bacteroidetes (2.79%), and Gemmatimonadetes (0.096%) (Figure 2A). Our method was effective at isolating members of the family Lachnospiraceae, with 542 total identified Lachnospiraceae isolates, representing the highest number of unique species (n = 73, 27.55%) (Figure 2B). Of these isolates, both *A. hadrus* (n = 139) were isolated multiple times across the nine total individuals, with at least one isolate of each retrieved from each individual, while *A. caccae* (n = 11) was isolated from only three individuals (Supplementary figure 1A). We observed that yeast casitone fatty acids (YCFA) media retrieved the highest number of total isolates (n = 249, 23.99%) and specifically Lachnospiraceae isolates (n = 149, 27.49%). However, Brain Heart Infusion (BHI) and BHI supplemented with rumen fluid (BHI+R) were also effective in targeting Lachnospiraceae species, with 139 isolates (25.65%) and 144 isolates (26.57%) respectively (Figure 2D). Increasing the number of total colonies picked per sample also improved recovery of diverse species (Supplementary Figure 1B).

**Figure 1.**
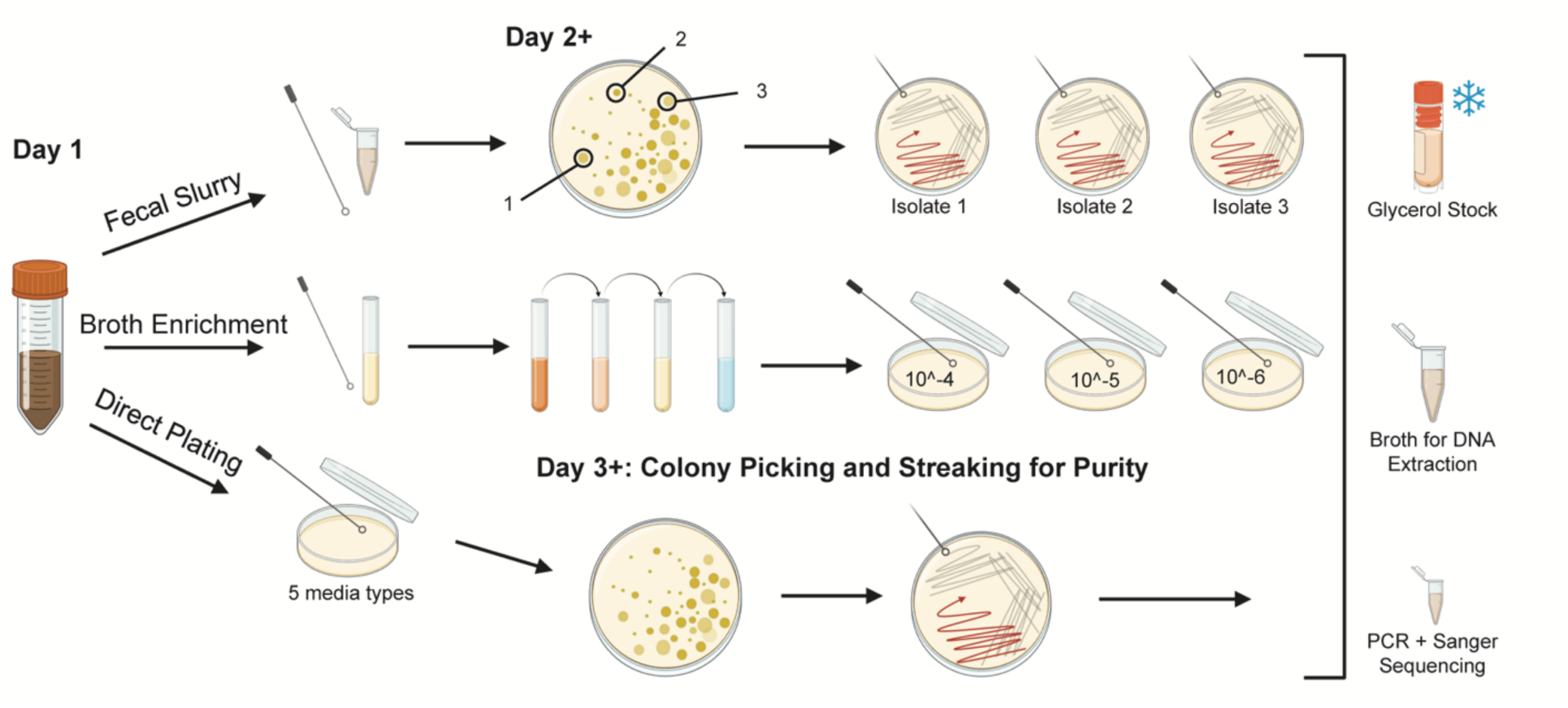
Isolation of human fecal bacteria. Graphical representation of the isolation method. Fecal samples (n=9) from healthy human were either streaked directly, streaked as a slurry, or enriched in liquid broth prior to dilution plating in different media types. Single, morphologically distinct colonies were streaked to purity and processed into glycerol stocks for long term storage and DNA extraction for Sanger and whole genome sequencing.

**Figure 2.**
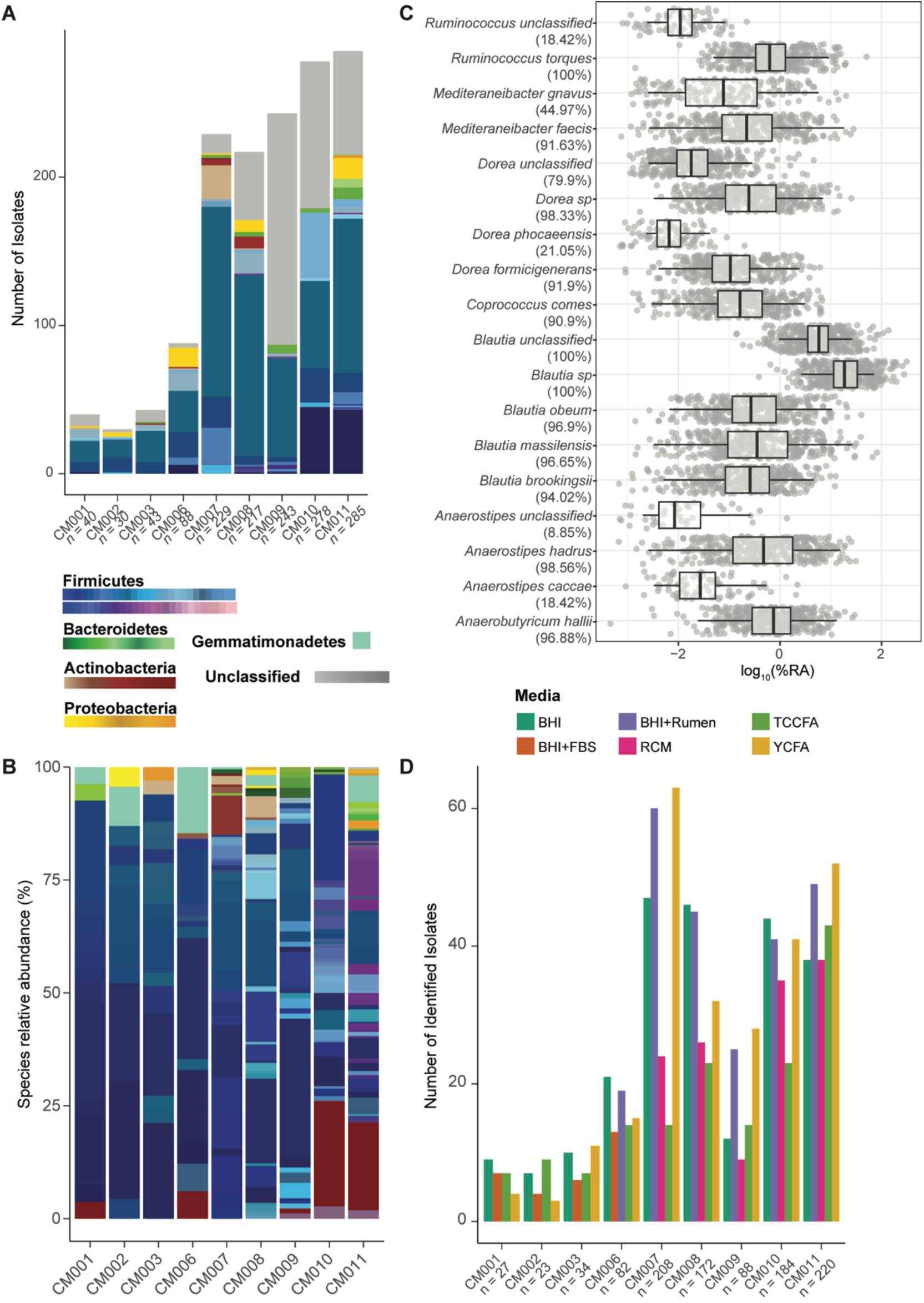
Lachnospiraceae are prevalent, abundant, and frequently isolated from human fecal samples. **A)** Total number of isolates obtained from the isolation protocol across nine human fecal samples, colored by taxonomic classification at the family level (n = number of colonies picked during isolation). **B)** Detection of select Lachnospiraceae species (using a custom classifier curated from isolated Lachnospiraceae species) in previously published 16S rRNA gene-based sequencing datasets from healthy human. Log_10_ of relative abundance of species is displayed on x-axis with prevalence (presence or absence of identified sequence). **C)** Relative abundance of unique species identified per sample, color-coded by genera. **D)** Total number of isolates identified by Sanger sequencing, grouped and color-coded by media type (n = number of colonies picked during isolation).

To broadly identify the prevalence of Lachnospiraceae species within the human gut, we created a custom classifier of the 16S rRNA gene from our sequenced isolates and mapped seven datasets of healthy adult fecal 16S rRNA amplicon sequences to these sequences (Figure 2B). As observed previously^70^, Lachnospiraceae species were highly prevalent, especially *Ruminococcus torques*, *Anaerostipes hadrus*, *Blautia obeum*, *Blautia massilensis*, and unclassified *Blautia* and *Dorea*, which were present in 95-100% of the samples from the datasets. Overall, relative abundance of most Lachnospiraceae species were low (∼0-25%), except for unclassified *Blautia* and *Blautia sp,* which exhibited up to 70% relative abundance for some samples. While *A. hadrus* was detected in 98% of the samples, *A. caccae* was only detected in ∼19% of the samples, concurring with the proportions of these species identified from our isolation pipeline.

### *Anaerostipes* genus is defined by 12 unique species

Given the prevalence of *A. caccae* and *A. hadrus* from our isolation pipeline, we sought to identify genomic diversity across the *Anaerostipes* genus. We compared whole genomes from a subset of our isolated *Anaerostipes* species (4 *A. caccae*; 17 *A. hadrus*) and 105 publicly available *Anaerostipes* genomes. An initial maximum likelihood tree based on single nucleotide polymorphisms (SNPs) demonstrated host-specific clustering with around 90% bootstrapping, supported by bootstrapping 500 times (Figure 3A, Supplementary Figure 2A). Genomes classified as *A. hadrus, A. amylophilus, and A. caccae* were exclusively found in humans, whereas *A. avistercoris, A. rhamnosivorans*, and *A. excrementavium* were exclusively from chickens. Furthermore, multiple genomically divergent *A. caccae* or *A. hadrus* strains were isolated from the same human sample in our isolation pipeline, suggesting coexistence of strains. For instance, the fecal sample from subject CM003 sample yielded three *A. hadrus* and two *A. caccae* strains. An additional UPGMA tree built from average amino acid identity (AAI) using EzAAI corroborated the SNP phylogeny (Supplementary Figure 2B).

**Figure 3.**
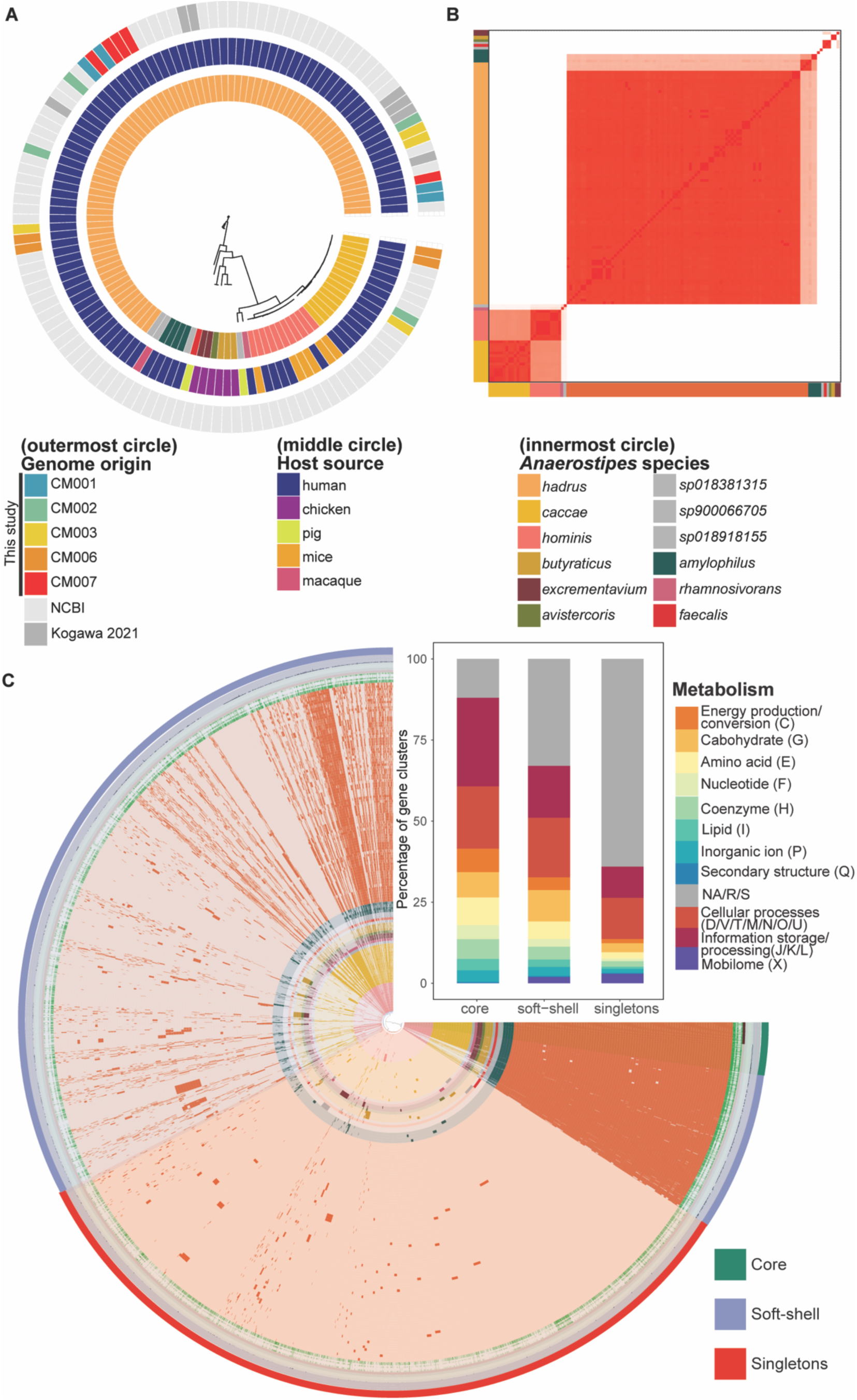
*Anaerostipes* genus is defined by 12 distinct species. **A)** Maximum likelihood tree based on single nucleotide polymorphisms (SNPs) in the core genome across 126 *Anaerostipes* genomes, overlayed with species, host source, and host subject (if isolated from this study) or published dataset. **B)** Average nucleotide identity (ANI) percentage across all 126 genomes. (Red denotes 100% ANI; white 80% ANI). **C)** Pangenome of *Anaerostipes* genus, as demonstrated by Anvi’o, displaying presence/absence of core (present in 100% of genomes), soft-shell (present in 99% of the genomes), and singletons (present only in a single genome across all genomes). Species color on the x and y axes are designated in legend. (Inset panel) Relative abundance of COG categories (color in legend) representing core, soft-shell, and singleton genes.

To identify additional clades within and across species, we dereplicated and calculated the average nucleotide identity (ANI) using Anvi’o. This identified 12 species within the *Anaerostipes* genus at < 95% ANI, with classification of these species matching their assigned taxonomy in GTDB (Figure 3B) which was further validated at <95% AAI (Supplementary Figure 2C). Overall, the pangenome of the genus *Anaerostipes* was open (Supplementary figure S3A), demonstrating a limited number of core genes (513 genes present in all 126 genomes), with most genes categorized as soft-shell core genes (12,138 genes present in at least two genomes) (Figure 3C). Most of the core genes within the genus were identified as related to amino acid synthesis and metabolism. Within amino acid production genes, most were specifically related to aromatic amino acid biosynthesis including 3-dehydroquinate synthase, which catalyzes the second step of Shikimate pathway^71^, shikimate kinase, fifth enzyme in the Shikimate pathway^72^, and aspartate aminotransferase, which catalyzes formation of oxaloacetate through the Krebs’ cycle^73^ (Figure 3C, inset panel). Most singleton genes (only present in one genome) were unknown (2630 of 4350 total singleton genes) as determined by the COG database. Of those identified, most genes belonged to transcription, defense mechanisms, and cell wall biogenesis categories of COG.

### *Anaerostipes* genus can be divided into three functional clusters

Functionally, the genomes split into three clusters, as determined by clustering using partition around medoids (PAM, average mean silhouette of the clusters = 0.91) of the principal coordinates of axes (PCoA) analysis calculated from a Bray-Curtis dissimilarity index of gene predictions using Prokka (Figure 4A). Cluster A (91 genomes) was dominated by *A. hadrus* genomes but also contained other human-associated genomes classified as *A. amylophilus*, *A. sp900066705* and *A. sp018918155*. Cluster B (28 genomes) was dominated by *A. caccae* and included the newly separated *A. hominis* and *A. sp018381315* genomes, from either human or other mammalian host origin. Cluster C (7 genomes) was dominated by species isolated from chickens (*A. butyraticus, A. avistercoris*, and *A. excrementavium*) and *A. faecalis* (isolated from pigs). Given the host specificity of the three clusters, we sought to identify common and unique functional features between the three clusters. Almost half of the identified annotated genes (46.4% of 2695 distinct genes total) were shared across all three clusters, with an average of 14 – 16% of genes unique to cluster A or B and 5% in cluster C (Figure 4B). Clusters A and B shared more genes (9.5%) compared to cluster C and A (6.2%) or C and B (3.8%). Of the shared genes that were identifiable within the COG database, most genes classified to functional category J: Translation, ribosomal structure and biogenesis (14%) followed by E: Amino acid transport and metabolism (10.7%) and G: Carbohydrate transport and metabolism (9.9%). Within the 120 COG identifiable features exclusive to cluster A and 111 features exclusive to cluster B, E: Amino acid transport and metabolism (10%) represented the largest functional category. However, for cluster C, which exhibited 33 exclusive features, G: Carbohydrate transport and metabolism (15%) was the major functional category (Supplementary Figure 3D).

**Figure 4.**
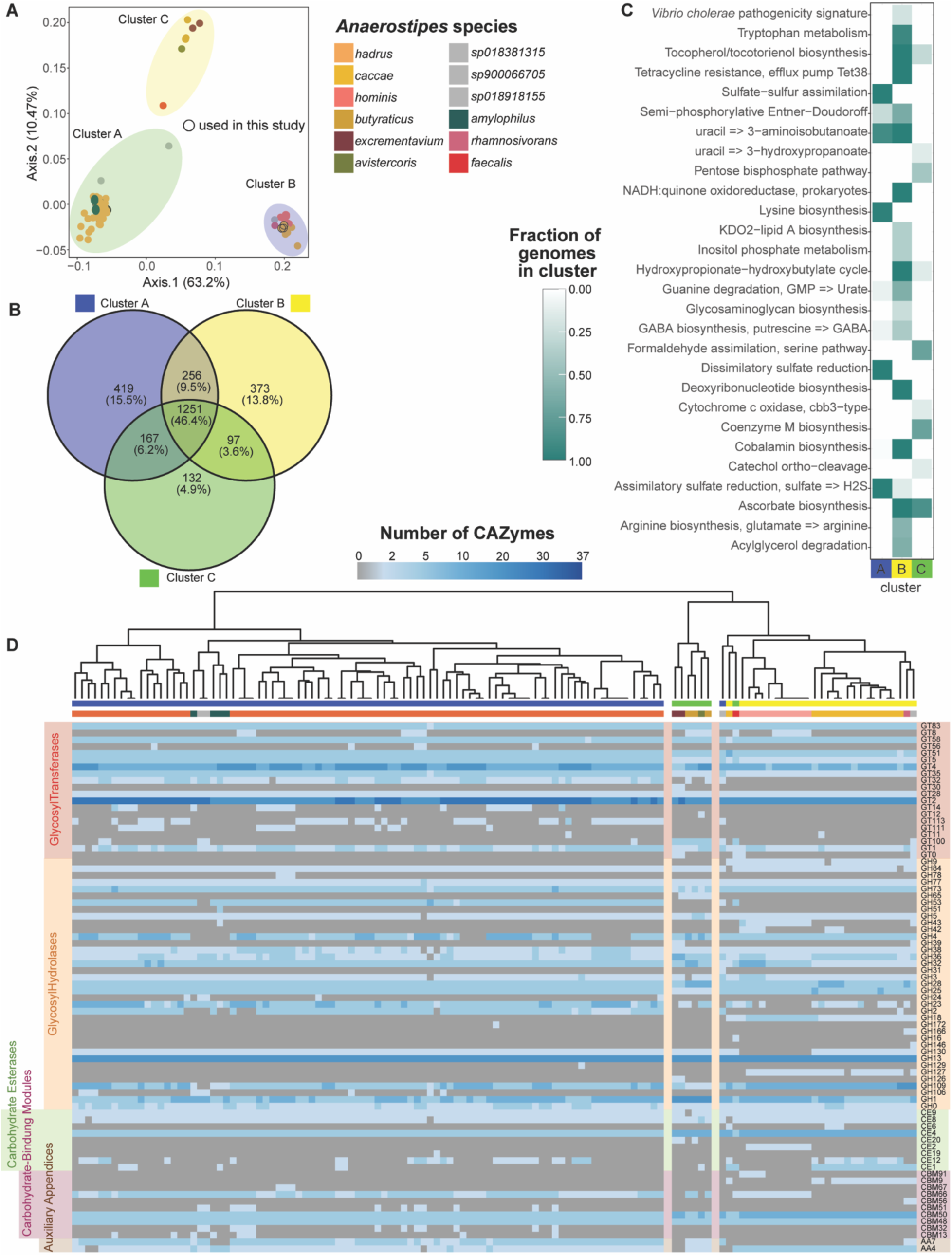
*Anaerostipes* clusters into three functional groups. **A)** Principal coordinates analysis (PCoA) based on a Bray-Curtis distance matrix of presence/absence of gene assignments generated using Prokka, colored by species. Clusters were assigned based on partitioning around medoids (PAM). **B)** Venn Diagram of predicted genes (Prokka) displaying total numbers and percentage of genes shared or unique to each cluster (n=2696 total classified genes). **C)** Differentially enriched KEGG modules across functional clusters (False Discovery Rate (FDR), q-value <0.05), colored by fraction of genomes within each cluster. **D)** Heatmap of number and type of carbohydrate-active-enzymes (CAZymes) in individual genomes (n = 126) using complete linkage clustering. GT = glycosyltransferase, GH = glycoside hydrolase, CE = carbohydrate esterase, CBM = carbohydrate-binding module, AA =Auxiliary appendices.

We also calculated functional enrichment across the clusters to identify differential representation of functional pathways (Figure 4C). Diverse functional pathways were overrepresented in cluster B, including pathways associated with amino acid metabolism (tryptophan metabolism; arginine biosynthesis) and nucleotide metabolism (guanine ribonucleotide degradation), with all genomes possessing a putative cobalamin biosynthesis pathway. All genomes in cluster A exclusively possessed pathways for sulfate assimilation or reduction, lysine biosynthesis, and pyrimidine degradation, whereas all cluster C exclusively genomes possessed a coenzyme M biosynthesis pathway. However, module completion (defined as > 75% complete) of many of the differentially identified pathways was only identified for lysine biosynthesis, one pyrimidine degradation pathway (uracil to 3-aminoisobutanoate), and sulfate assimilation (Supplementary Figure 4).

Variation of carbohydrate-active enzymes (CAZymes) were observed across the genomes, which closely matched clustering based on overall gene content above (Figure 4D). The glycosyltransferase category GT2 was the most numerous categories of CAZymes observed across all genomes, although Cluster A genomes demonstrated increased diversity of glycosyltransferases. Conversely, the diversity of carbohydrate esterases and carbohydrate-binding modules was higher in clusters B and C, suggesting an affinity to substrates different from cluster A, an indication of nutrient niches specific to the respective clusters.

### Distribution of genes associated cobalamin and short chain fatty acid production varies across *Anaerostipes* functional clusters

We next sought to identify how genes previously associated with select *Anaerostipes* species were distributed across all genomes. Corrinoids, such as cobalamin or B12, are important for gut microbial physiology since they are an essential co-factor for corrinoid dependent enzymes^74^. Production of cobalamin by *A. caccae* has been postulated, as co-cultures of *Akkermansia mucinphila* and *A. caccae* have been demonstrated to produce low propionate levels through corrinoid-dependent methylmalonyl-CoA mutase enzymes in *A. muciniphila*^75^. We observed that all species in cluster A (containing *A. caccae*) possessed genes belonging to the main, aerobic, anaerobic, C4 and salvage pathways to produce cobalamin (Figure 5A). While only the salvage pathway represented by *cobU* and *btuR* in cluster B genomes was present, both the main and aerobic pathways were complete or near complete, only missing the presence of the final genes *pduO* (adenosyltransferase) and *cobST* (cobalamin-5’-phosphate synthase) in some strains, suggesting modified corrinoid production. The anaerobic pathway in most genomes lacked *cbiK* (anaerobic cobalamin biosynthetic cobalt chelatase), which may corroborate previous co-culture observations requiring the presence of additional bacteria to produce cobalamin^75^. In contrast, clusters A and C lacked almost all cobalamin-related genes, with the exception of the *sp018918155* genome in cluster A.

**Figure 5.**
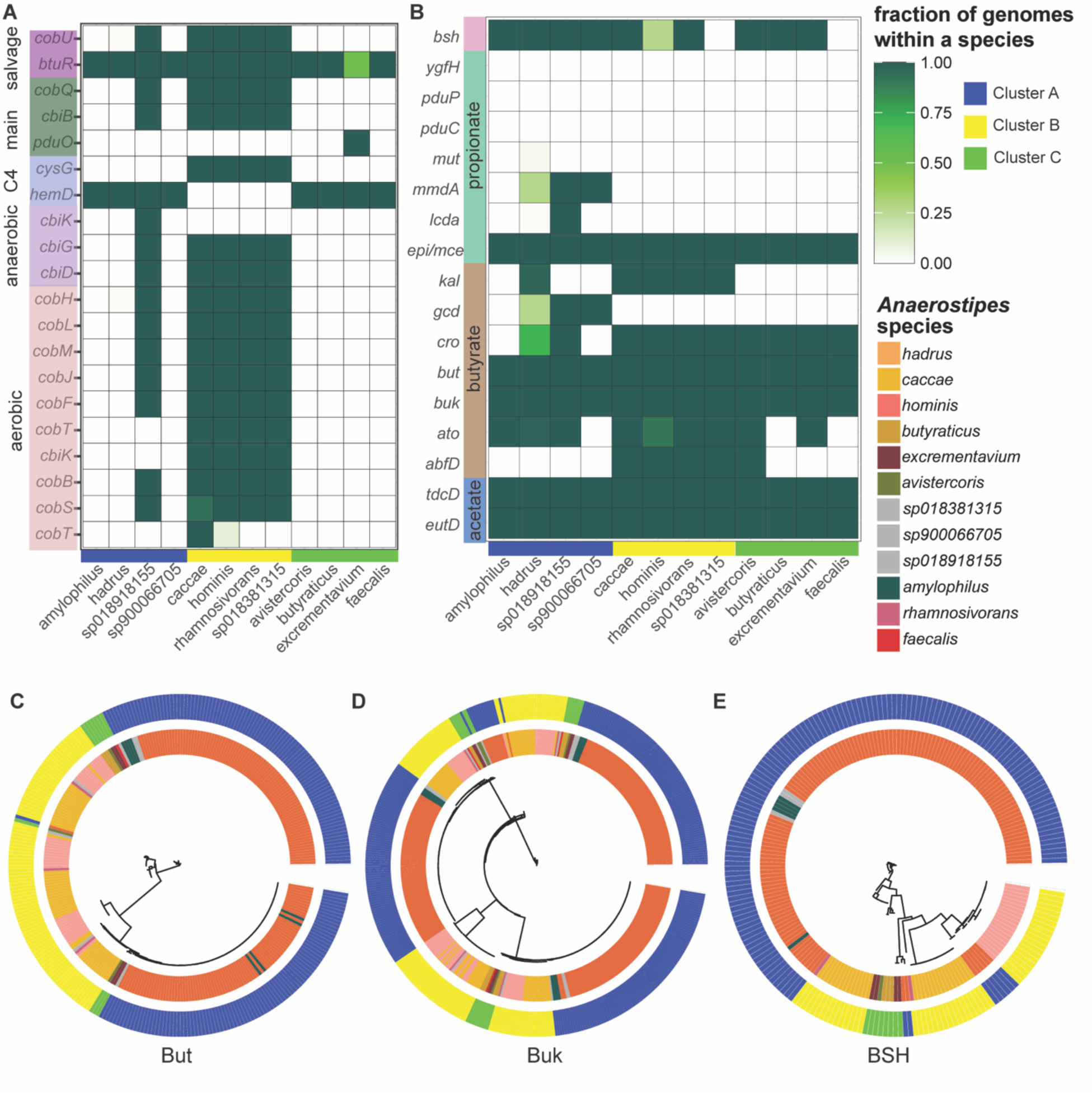
*Anaerostipes* species display varied capacity for cobalamin production, short chain fatty acid (SCFA) production, and bile salt hydrolases (BSH). **A)** Presence or absence of selected genes from aerobic, anaerobic, main (cob(II)yrinic acid a,c-diamide to adenosylcobalamin) salvage, and C4 cobalamin production pathways represented as a fraction of genomes within a species. **B)** Presence or absence of selected genes from acetate, butyrate, and propionate pathways and BSH, represented as a fraction of genomes within each species. **C)** Maximum likelihood tree generated from butyrate CoA: acetate CoA transferase (But) protein hits (identity > 60%) in all genomes (bootstrap n=100) **D)** Maximum likelihood tree generated from butyrate kinase (Buk) protein hits (identity > 60%) in all genomes (bootstrap n=100) **E)** Maximum likelihood tree generated from BSH protein hits (identity > 60%) in the genomes (bootstrap n=100).

A major health-related function attributed to *Anaerostipes* is the production of SCFAs, especially butyrate^23^. We observed widespread presence of butyrate- and acetate-related genes across all species and functional clusters, with some variability associated with the functional clusters (Figure 5B). Importantly, the main butyrate-associated genes *but* (butyryl-CoA:acetate CoA transferase) and *buk* (butyrate kinase), both terminal enzymes that catalyze production of butyrate, were present in all genomes. A maximum likelihood tree of the aligned amino acid sequences represented by *but* and *buk* demonstrated species-specific clustering with low bootstrap values (< 50%, n = 500), suggesting near identical sequences across all genomes (Figure 5C, D). The presence of other SCFA genes were more specific to functional clusters. Most cluster B and C genomes lacked *gcd* (glutaconyl-CoA decarboxylase), that catalyzes conversion of glutaconyl-CoA to crotonyl-CoA via the glutarate pathway of butyrate biosynthesis, and cluster C genomes lacked *kal* (3-aminobutyryl-CoA ammonia lyase), which catalyzes conversion of 3-aminobutyryl-CoA to crotonyl-CoA. In contrast, only *epi/mce* (methylmalonyl-CoA epimerase), which converts succinate to propionate, from the propionate pathway was present across all genomes^76^. Some cluster A genomes exhibited the presence of *mmdA* (methylmalonyl-CoA decarboxylase), which catalyzes S-methylmalonyl-CoA to propionyl-CoA, and *lcdA* (lactoyl-CoA dehydratase), which converts lactoyl-CoA to acryloyl-CoA^77^.

An important function of some commensal gut microbes is deconjugation of primary bile acids, performed through bile salt hydrolases (*bsh*). All species demonstrated the presence of *bsh*, with the exception of *A. faecalis* and *sp018381315*, which were only represented by one genome each, and a few *A. hominis* genomes (Figure 5B). Although bootstrap values for amino acid sequence comparison represented by BSH were low, some sequences were more closely related to each other than others, such as AGGJHBAO_01892 from CM03_47 (*A. hadrus*), OMENAAMB_02178 from CM02_14 (*A. hadrus*) and EGPMHLCL_02247 from CM03_84 (*A. hadrus*), for which bootstrapping values were at 100%, even though all three strains were isolated from two different human subjects (Figure 5E).

### Pangenomic comparisons of *A. caccae*, *A. hominis* and *A. hadrus* demonstrate functional differences

We next focused on comparing functional differences within and across three prevalent human-associated *Anaerostipes* species. Pangenomic comparisons within *A. hadrus* (84 genomes), *A. caccae* (15 genomes), and *A. hominis* (11 genomes) species suggested that both *A. hadrus* and *A. caccae* displayed open pangenomes (alpha < 1), whereas *A. hominis* demonstrated a closed pangenome (alpha > 1) (Figure 6A). Based on identified COG genes, *A. hadrus* demonstrated the smallest core genome among the three species, with conservation of J: Translation, ribosomal structure and biogenesis (representing 10.69%) across the *A. hadrus* core genome while maximum genes belonged to G: Carbohydrate transport and metabolism genes (8.75%) and X: Mobilome genes (3.9%) in soft-shell and singleton categories, respectively (Supplementary Figure S5). This was in comparison to *A. caccae* and *A. hominis* core genes, where G: Carbohydrate transport and metabolism comprised 12.7% and 13.1% of core genes, respectively. *A caccae* and *A. hominis* also exhibited a smaller percentage of X: Mobilome genes (0.14% and 0.61%, respectively) (Supplementary figure S5).

**Figure 6.**
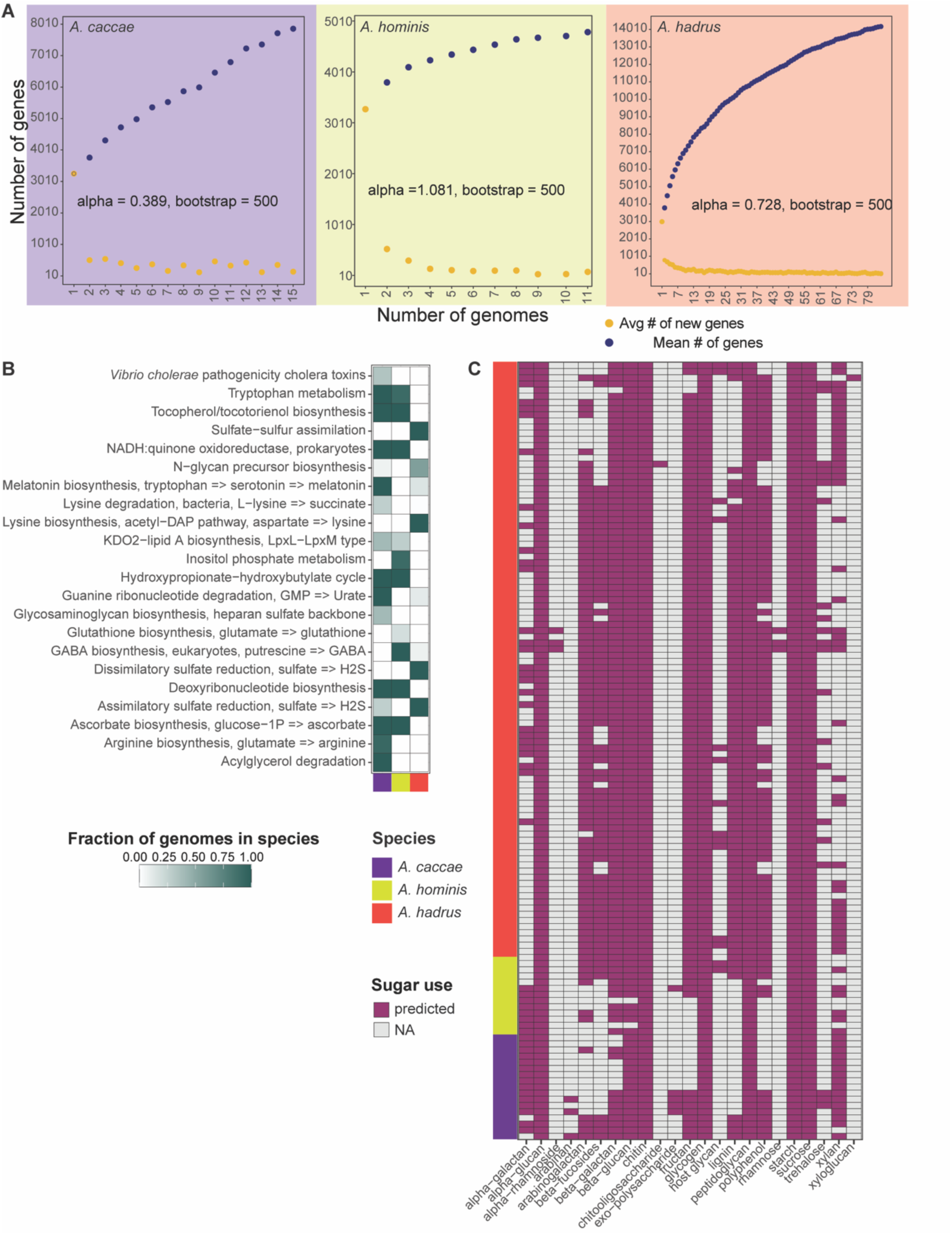
Human-associated *Anaerostipes* species, *A. hadrus*, *A. caccae* and *A. hominis*, vary in functional capacities. **A)** Number of genes as a function of number of genomes depicting number of new genes in yellow dots and number of genes in the pangenome in blue dots of *A. caccae* (n=15, purple), *A. hominis* (n=11, seagreen) and *A. hadrus* (n=84, red). Alpha equals the power law estimate, ran over 500 iterations using micropan in R (p-value < 2e^-16). **B)** Differentially enriched KEGG modules across the three species (False Discovery Rate (FDR), q-value <0.05), colored by fraction of genomes. **C)** Presence (purple box) or absence (grey box) of sugar substrate (x-axis) predicted from carbohydrate-active enzymes (CAZymes) across all the genomes in the 3 species.

To investigate functional differences across the three species, we used Anvi’o to identify differentially enriched modules. *A. caccae* and *A. hominis* shared more similarities compared to *A. hadrus*, including exhibiting pathways associated with tryptophan metabolism, tocopherol/tocotorienol biosynthesis, and NADH:quinone oxidoreductase (Figure 6B). Melatonin and arginine biosynthesis, as well as guanine ribonucleotide and acylglycerol degradation were almost exclusively observed in *A. caccae*, whereas *A. hominis* genomes exhibited inositol phosphate metabolism and GABA biosynthesis pathways. In comparison, *A. hadrus* demonstrated widespread presence of multiple pathways associated with sulfate reduction and lysine biosynthesis.

We also used dbCAN3 to predict the substrate carbohydrate use across all genomes (Figure 6C). Several substrate predictions were conserved across all three species, including alpha-glucan, starch and sucrose, despite genomic variation observed within the species. Arabinan, xyloglucan, and exopolysaccharides were not predicted to be metabolized by these three species. Use of lignin and beta-fucosides seemed restricted to *A. hadrus*, and *A. caccae* genomes generally lacked CAZymes involved in fructan and polyphenol utilization, suggesting that *A. caccae* may exhibit a more restricted carbohydrate use profile. Comparison of individual CAZymes followed this pattern, whereby *A. caccae* demonstrated a less diverse CAZyme profile (Figure 4D).

### *Anaerostipes* species exhibit variation in factors associated with colonization

While 16S rRNA gene-based studies commonly associate *Anaerostipes* with health, some studies have identified associations to particular disease states^78,79^. We screened all *A. caccae*, *A. hadrus*, and *A. hominis* genomes against virulence and antibiotic resistance gene databases using PathoFact^67^. All genomes demonstrated the presence of toxin-antitoxin systems, which may represent conserved plasmid and transposon maintenance (Supplementary figure S6). All *A. caccae* and *A. hominis* genomes exhibited the presence of exfoliative toxins A and B, initially identified on *Staphylococcus aureus*^80^. Some strains of *A. caccae* also exhibited the presence of a gene encoding a zona occludens toxin, an enterotoxin located on prophages of some pathogenic *Vibrio* species^81^. Most *A. hadrus* and *A. hominis* genomes also exhibited the presence of a gene encoding an insecticidal toxin complex protein, TccC that has been associated with increased colonization ability of some pathogenic *Yersinia* strains^82,83^.

We also used PathoFact to investigate the putative antimicrobial profiles across the 3 major *Anaerostipes* species. All genomes contained widespread glycopeptide, macrolides, lincosamides, and streptogramines (MLS), and efflux pump encoding multidrug resistance genes, which can be attributed to either accumulation of mutations on exposure or dissemination of resistance through horizontal gene transfer across the gut microbial community^84^ (Figure 7A). *A. hominis* genomes and some strains of *A. hadrus* exclusively exhibited genes encoding beta-lactam and polymyxin resistance, which were absent in all *A. caccae* genomes. Genes associated with resistance to other antibiotics, such as tetracycline, aminoglycosides, and bacitracin, were observed sporadically throughout the 3 species.

**Figure 7.**
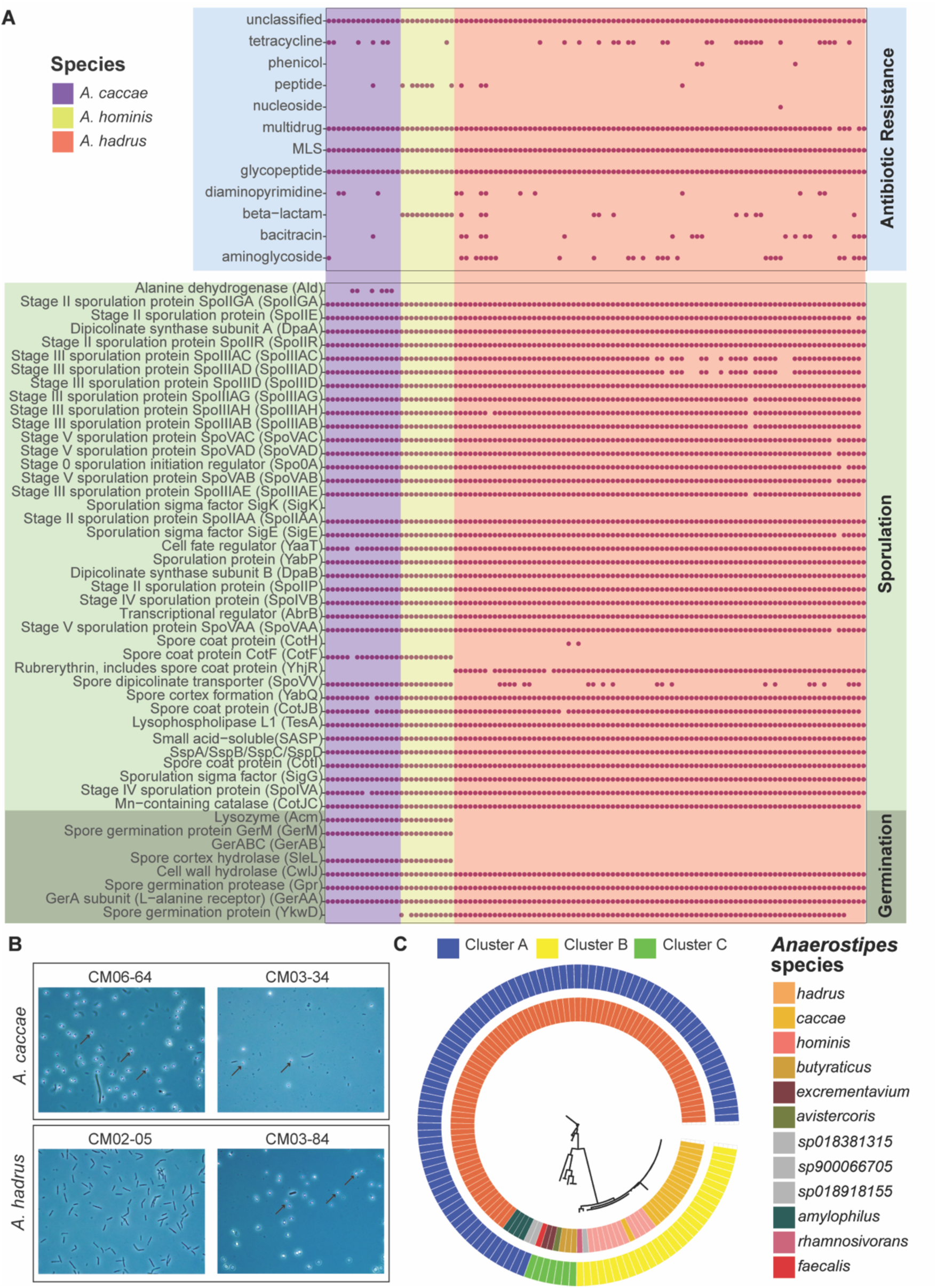
*A. caccae* and A. hadrus isolates display common antibiotic and sporulation. **A)** Presence (purple dot) or absence (no dot) of antibiotic resistance, sporulation, and germination genes across *A. caccae* (n = 15), *A. hominis* (n = 11) and *A. hadrus* (n = 84) genomes. Antibiotic resistance was detected through PathoFact and sporulation or germination genes through KOfam. **B)** Phase contrast of spores from two *A. caccae* and two *A. hadrus* strains isolated in this study. Spores marked by arrows (no spores detected in CM02-05 strain). **C)** Maximum likelihood tree for Spo0A protein hits (identity>60%) across all 126 genomes (bootstrap n=100).

Finally, we were interested in identifying potential sporulation and germination pathways across *A. hadrus*, *A. hominis*, and *A. caccae*. The formation of endospores across various *Anaerostipes* species has been observed but not fully characterized^21^. All three species exhibited the presence of most genes required to sporulate through the mechanism characterized for *B. subtilis*^85^, including *spo0A* (sporulation initiation protein), stage II, stage III and stage V sporulation proteins, and various spore coat proteins (Figure 7A). Some species-specific conservation was observed, such as the presence of *ald* (alanine dehydrogenase) exclusively in *A. caccae* genomes and *yhjR* (rubrerythrin, spore coat) exclusively in *A. hadrus.* Given the previously observed variability in *Anaerostipes* spore formation^21^, we tested the ability of two *A. caccae* and two *A. hadrus* strains to form spores *in vitro*. Both *A. caccae* strains formed visible spores after fourteen days of growth in Clospore media, while only one *A. hadrus* strain produced visible spores under phase contrast microscopy (Figure 7B).

We observed limited variation within sporulation and germination genes between *A. hadrus* and *A. caccae* strains. However, on closer inspection of the genomes, we found that the two tested *A. caccae* strains, CM03-34 and CM06-64, did not have *ykwD* (PDB:1CFE) and *yhjR* (rubrerythrin, includes spore coat protein) (PDB:2E0Z) related genes, while the two tested *A. hadrus* strains did not possess genes for *cotF* (spore coat protein), *acm* (lyzozyme M1 (1,4-beta-N-acetylmuramidase), GH25 family) (PDB:1H09), *sleL* (spore cortex hydrolase, GH18) (PDB:4S3J), *gerM* (spore germination protein) (PDB:6GZB) and *spoVV* (spore dipicolinate transporter) based on the COG database. We also observed genomic differences in the sporulation and germination genes between the two tested *A. hadrus* strains. These differences were mostly observed in sporulation genes, such as *spoIVFB*, *spoIIP*, *yabP*, *spoVAD*, and *spoIIID*, and CM02-5 did not seem to possess *spoVAC* at all, which is an important germination related mechanosensitive channel in *B. subtilis*^86^.

Given the conservation of *gerA* across these species, we further sought to compare *gerAA*, *gerAB*, and *gerAC* (the three genes encoding each of the three proteins involved in a functional GerA receptor)^87^ from COG profiles. Comparison to the COG database identified only one *gerA* protein, GerAA, in all three genomes, but no GerAB in any genomes. In contrast, use of a UniProt database using DIAMOND identified presence of *gerAB* homologues, which encodes for the protein that typically binds to the germinant signal, but no *gerAA,* which encodes for the protein that transduces a downstream signal^87^., suggesting a discrepancy in database identification. The *gerAC* gene, which encodes for a lipoprotein with an unknown function^88^, was not detected in any species. Aligning the GerAB protein sequence using DIAMOND revealed tight clustering by species (bootstrapping > 50%), suggesting species-specific germination pathways (Figure 7C).

## DISCUSSION

Our study aimed to genomically characterize species within *Anaerostipes*, a genus known to inhabit the human gut microbiome. *Anaerostipes* species have been identified across global human populations^89^, as well as other animal hosts^90^. Using both cultivation and 16S rRNA comparisons, we observed high prevalence of *Anaerostipes* species in the human gut, particularly for the species *A. hadrus*. *Anaerostipes* genomes could be divided into three main functional clusters. Apart from SCFA production, certain *Anaerostipes* species could potentially contribute to other important secondary metabolites, such as Vitamin B12. These data, alongside multiple studies correlating *Anaerostipes* with health^78,79,91,92^ and any obvious virulence, support a commensal (and thus a potentially important) role of *Anaerostipes* in the human gut^93^.

Isolation of multiple *Anaerostipes* species revealed that multiple ‘strains’ (defined as phylogenetic clusters with a 99% ANI within a species in our study) were present in a single individual. For example, fecal sample from subject CM003 yielded three *A. hadrus* strains and one *A. caccae* strain. This suggests co-existence between different strains of the same species within the same host and existence of strains of different species of *Anaerostipes* within the same host, all indicating niche specialization within a host. Similar results have been reported for *E. coli*, *R. gnavus* and *B. vulgatus*, where multiple strains of the same bacteria have been identified within the same individual^94^. We also observed the existence of two highly similar strains of *A. caccae* (dereplicating at 99% ANI) across different individuals (CM002 and CM003), demonstrating that specific *Anaerostipes* species may be generalizable across the human gut.

The genomic comparisons across our isolates and available genomes suggests that *Anaerostipes* display high intra-species similarity. Our phylogenetic, AAI, and ANI comparisons matched the reported taxonomic identification of available genomes included in our study, revealing 12 species in total, including three unnamed species. Some of the named and unnamed species were represented by single genomes only due to quality control. Phylogenetically, these species aligned with host specificity, with species such as *A. avistercoris* and *A. excrementavium* originating exclusively from chickens and *A. hadrus* and *A. caccae* originating from humans. This suggests adaptation of particular *Anaerostipes* species to their respective host environment, similar to co-speciation observations made for other gut symbionts such as *Bifidobacterium* and *Enterococcus* observed across host environments^95,96^. A previous study compared 527 assembled genomes from isolates and metagenome assembled genomes (MAGs) and found three main phylogenetic clusters within *A. hadrus* with no particular association between geographical regions^89^. Our study incorporated any genomes from this previous study that passed our quality control metrics (n = 41). We did not observe large-scale functional clustering within *A. hadrus*, but did observe 39 fairly homologous clusters at 99% ANI (obtained from Anvi’o). We also did not observe clustering within *A. hadrus* genomes using AAI (obtained from EzAAI) either. One explanation for this may be our smaller sample size, given our focus on high-quality genomes (which eliminated nearly all metagenome-assembled genomes).

Our comparisons across the 12 *Anaerostipes* species revealed potential host-specific speciation. While cluster A (*A. hadrus*-dominated) was almost exclusively described by human-sourced isolates, cluster C included genomes sourced from chickens and pig, whereas cluster B consisted of genomes sourced from humans, mice, and pigs. The impact of host physiology and phylogeny on the gut microbiome has been characterized in non-human primates, wherein specific gut microbiome signatures have been associated with host species despite similar dietary niches of the hosts^97^. Convergent evolution of gut bacterial species to perform similar functions in different hosts, while typically overlooked if only assessing taxonomy or phylogenetic similarity, has been observed across other hosts, such as within speciation of Enterobacteriaceae species across different *Ceanorhabditis* species, *C. elegans* and *C. briggsae*^98^.

Both shared and differentiated features across the three identified functional categories suggest independent roles or adaptation for *Anaerostipes* species in the gut. All species exhibited the canonical *but* (butyryl-CoA:acetate CoA transferase) and *buk* (butyrate kinase) genes that have defined *Anaerostipes* as a butyrate-producing genera. Additionally, all species exhibited the presence of *tdcD* (acetate propionate kinase) and *eutD* (phosphotransacetylase), associated with acetate production^99^, and *epi/mce* (methylmalonyl-CoA epimerase), associated with propionate production^100^. Almost all species exhibited the presence of *bsh* (bile salt hydrolase), adding to the growing list of bacteria capable of modifying host-produced primary bile acids^101^. However, differences across the functional clusters, which also displayed host specificity, may further distinguish species niches within the gut. Cluster B genomes (*A. caccae*-dominated) exhibited pathways for tryptophan metabolism, as well as other biosynthetic pathways to produce arginine, tocopherol (vitamin E), and cobalamin (vitamin B12). Closer comparison of cobalamin production revealed that *A. caccae* and other functionally similar species exhibited multiple cobalamin biosynthetic pathways, which has not been previously described in *Anaerostipes* specifically^102,103^. Although actual cobalamin production would have to be experimentally validated, the near completeness of these pathways suggests that *A. caccae* may play at least a cross-feeding role in cobalamin production in the gut^75^. Species within cluster A exhibited more diverse butyrate-production pathways compared to clusters B or C, although clusters A (*A. hadrus*-dominated) and B (*A. caccae*-dominated) exhibited additional genes, such as *kal* (3-aminobutyryl-CoA ammonia lyase), and *cro* (crotonase/enoyl-CoA hydratase). Comparison of But, Buk and BSH protein sequences across all species indicated conserved protein sequences through low bootstrap/branch support values.

Further investigation of the three most dominant human *Anaerostipes* species (*A. hadrus*, *A. caccae*, and *A. hominis*) revealed additional information relevant to human health. Both *A. caccae* and *A. hadrus* exhibited open pangenomes, a feature of sympatric bacteria living in close ecological contact with each other and suggesting additional gene diversification is possible should larger-scale comparisons within the species be conducted^104^. In contrast, *A. hominis* exhibited a closed pangenome, a characteristic of allopatric lifestyle, where organisms living in isolated environments have limited access to external genetic materials^105,106^. While it is possible that *A. hominis* has a very specialized niche and function in both mice and humans, it was not heavily represented in our analyses (11 genomes); additional genomes could reveal more novel genes, particularly given the presence of mobilome genes in the singleton pangenome bin. Both *A. caccae* and *A. hominis* also displayed the presence of more putative virulence genes compared to *A. hadrus*, including homologs for a *Vibrio* zona occludens toxin^107^ and *Staphylococcus* exfoliative toxins^108^. Despite some studies associating *A. caccae* with disease states rather than health^109^, there is no evidence of exfoliative toxins in gut microbes in current literature which could indicate a novel alternative function of the toxin structure, or a limitation of the database used. In terms of antibiotic resistance, *A. hadrus*, *A. caccae* and *A. hominis* had diversity in the presence of antibiotic resistance genes within and among the species, with *A. caccae* not possessing beta-lactam and polymyxin resistance genes at all.

Understanding sporulation and germination mechanisms of gut anaerobes is perhaps most relevant to development of microbial therapeutics, given that spore delivery is an optimal method to deliver anaerobic bacteria to the gut^110^. Although spore formation is encompassed by many commensal Clostridia and specifically Lachnospiraceae^111^, sporulation and germination mechanisms are incompletely described for species other than *Bacillus subtlitis* and the gut pathogen, *Clostridioides difficile*^112,113,114^. All three major human *Anaerostipes* species possessed sporulation and germination associated genes observed in *B. subtilis*, which sporulates via Spo0A phosphorylation and germinates through Ger proteins with GerAB that bind to the germinant nutrient L-alanine^112,113^. Initial comparisons using the COG database identified GerAA in the three *Anaerostipes*; however, closer phylogenetic analysis revealed that this was likely GerAB instead. We also observed a species-specific clustering of GerAB protein sequence informed by the bootstrapped maximum likelihood tree. This suggests *Anaerostipes* species might follow a germination mechanism similar to *B. subtilis*. While a previous study by Kodawaki et al. were not able to identify *in vitro* production of spores despite the identification of similar spore-related genes in *A. hadrus*^21^, we observed *in vitro* spore production of both tested *A. caccae* strains and one *A. hadrus* strain. We also observed a few genes such as *spoII* absent from CM02-5, the strain that did not produce spores. This suggests possible strain-to-strain variation across *A. hadrus* sporulation, warranting additional validation and comparison using lab-based methods.

Overall, our study aimed to define *Anaerostipes* across host species, elucidating genomic strain diversity that might influence previous associations based on 16S rRNA gene-based analyses alone. Our data suggests that while all *Anaerostipes* display widespread SCFA capability, certain *Anaerostipes* species may be host-restricted, demonstrating metabolic differences that may explain their adaptation to particular host gut niches.

## ACKNOWLEDGEMENTS

We would like to thank the participants who consented to donating fecal samples. This work was supported by grant number R35GM150609 (AMS). We would also like to acknowledge Clemson University for generous allotment of compute time on Palmetto cluster. This material is based on work supported by the National Science Foundation under Grant Nos. MRI# 2024205, MRI# 1725573, and CRI# 2010270. This publication was made possible, in part, with support from the Clemson University Genomics and Bioinformatics Facility, which receives support from the College of Science and two Institutional Development Awards (IDeA) from the National Institute of General Medical Sciences of the National Institutes of Health under grant numbers P20GM146584 and P20GM139769.

D.B. - Conceptualization, Data Curation, Formal analysis, Investigation, Methodology, Software, Project Administration, Writing—original draft, and Writing—review and editing; L.M. – Data Curation, Investigation, Methodology, Writing – review and editing; M.L.S. – Methodology, Investigation, Writing – original draft, Writing – review and editing; A.M.S. – Conceptualization, Project Administration, Supervision, Funding Acquisition, Writing—original draft, and Writing – review and editing.

