## Supplementary Figures for "Diversity and prevalence of *Anaerostipes* in the human gut microbiota"

### Supplementary Figures with Figure Legends

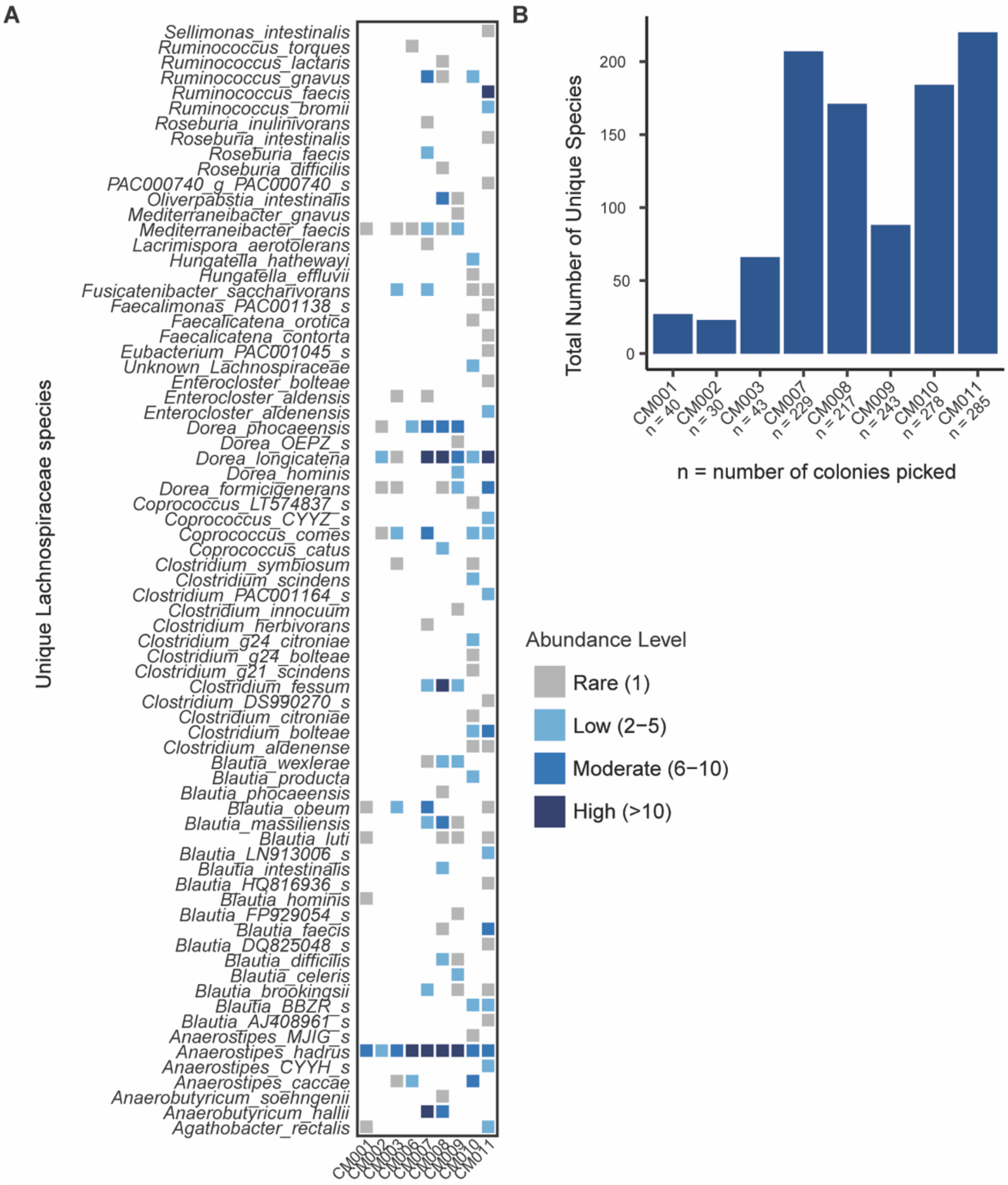


**Figure S1. Rescue of unique species increases with the number of colonies picked. A)** Number of strains (abundance) of unique Lachnospiraceae species isolated from human fecal samples (subjects on the y-axis). White boxes indicate absence of the strain. **B)** Total number of unique species, across nine human fecal samples (n = number of colonies picked during isolation).

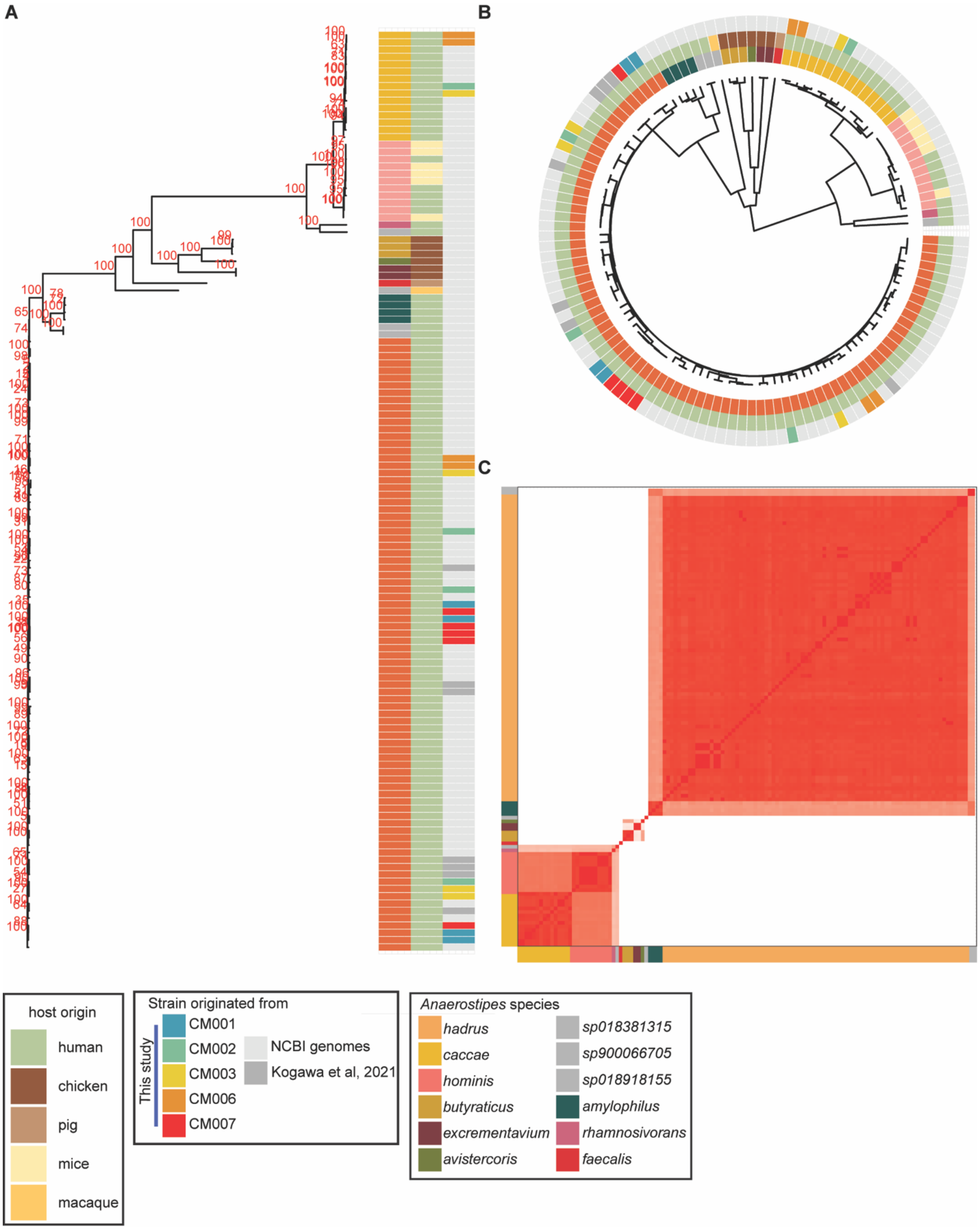


**Figure S2. Maximum likelihood trees of *Anaerostipes* based on single nucleotide polymorphisms (SNPs) in the core genome and Average amino acid identity (AAI). A)** Maximum likelihood tree made from SNPs in the core genome with bootstrap values in red text overlayed with species color, host source, and subject. **B)** UPGMA tree of Anaerostipes based on amino acid identity made using ezAAI overlayed with species color, host source, and subject. **C)** Average amino acid identity (AAI) percentage across all 126 genomes, colored by species. (Red denotes 100% ANI; white 80% ANI).

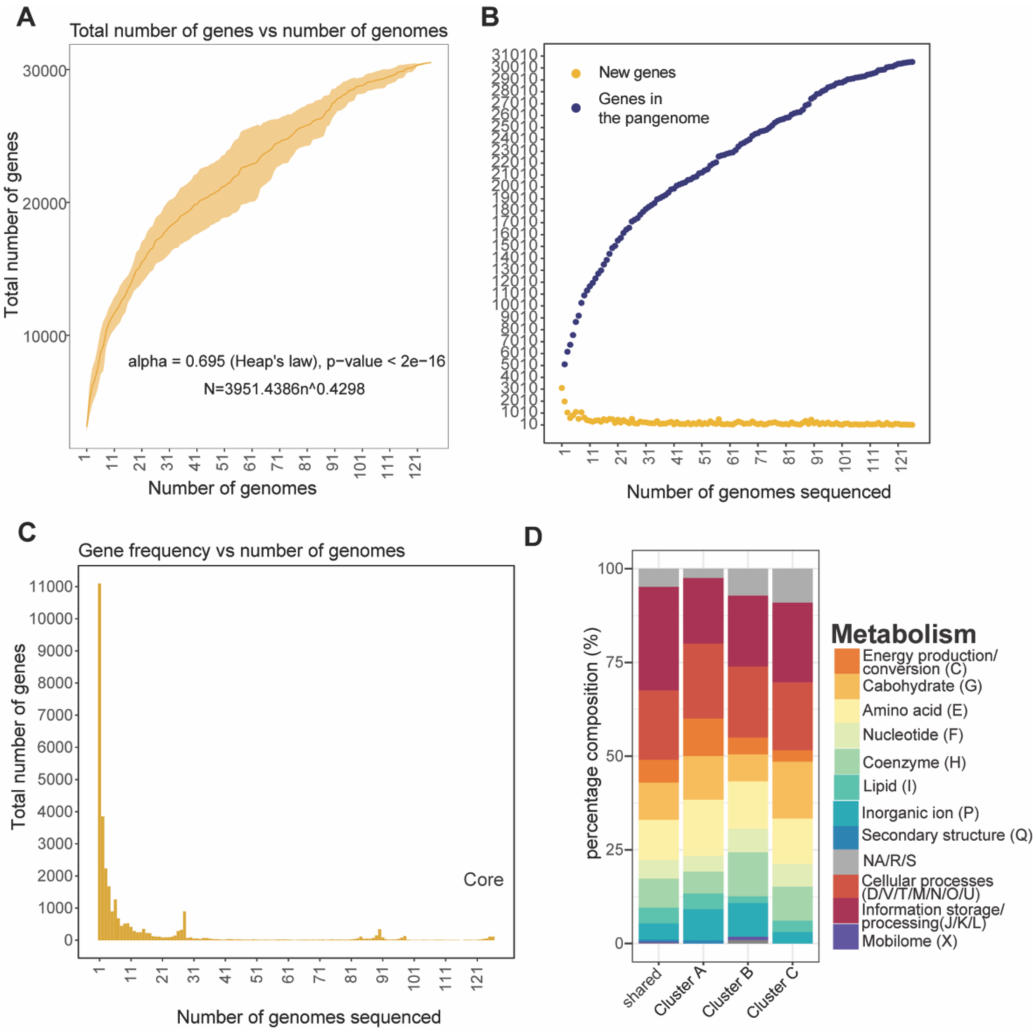


**Figure S3.** ***Anaerostipes*** **pangenome statistics. A)** Total number of genes as a function of number of all included genomes (n =126). Yellow colored solid line represents the average number of total genes from five subsamplings, with transparent yellow as error bars. **B)** Number of genes as a function of number of genomes depicting number of new genes in yellow dots and number of genes in the pangenome in blue dots of all genomes (n=126). Yellow colored boxplots represent the average number of total genes from five subsamplings. **C)** Total number of unique genes as a function of number of genomes (p-value < 2e^-16). Core genes depicted at 126 genomes by "Core". Alpha = Heap's law estimate, ran over 500 iterations using micropan in R (p-value < 2e^-16). **D)** Relative abundance of COG categories for genes (n=1251) common to each cluster, genes unique to cluster A (n=419), cluster B (n=373), and cluster C (n=132).

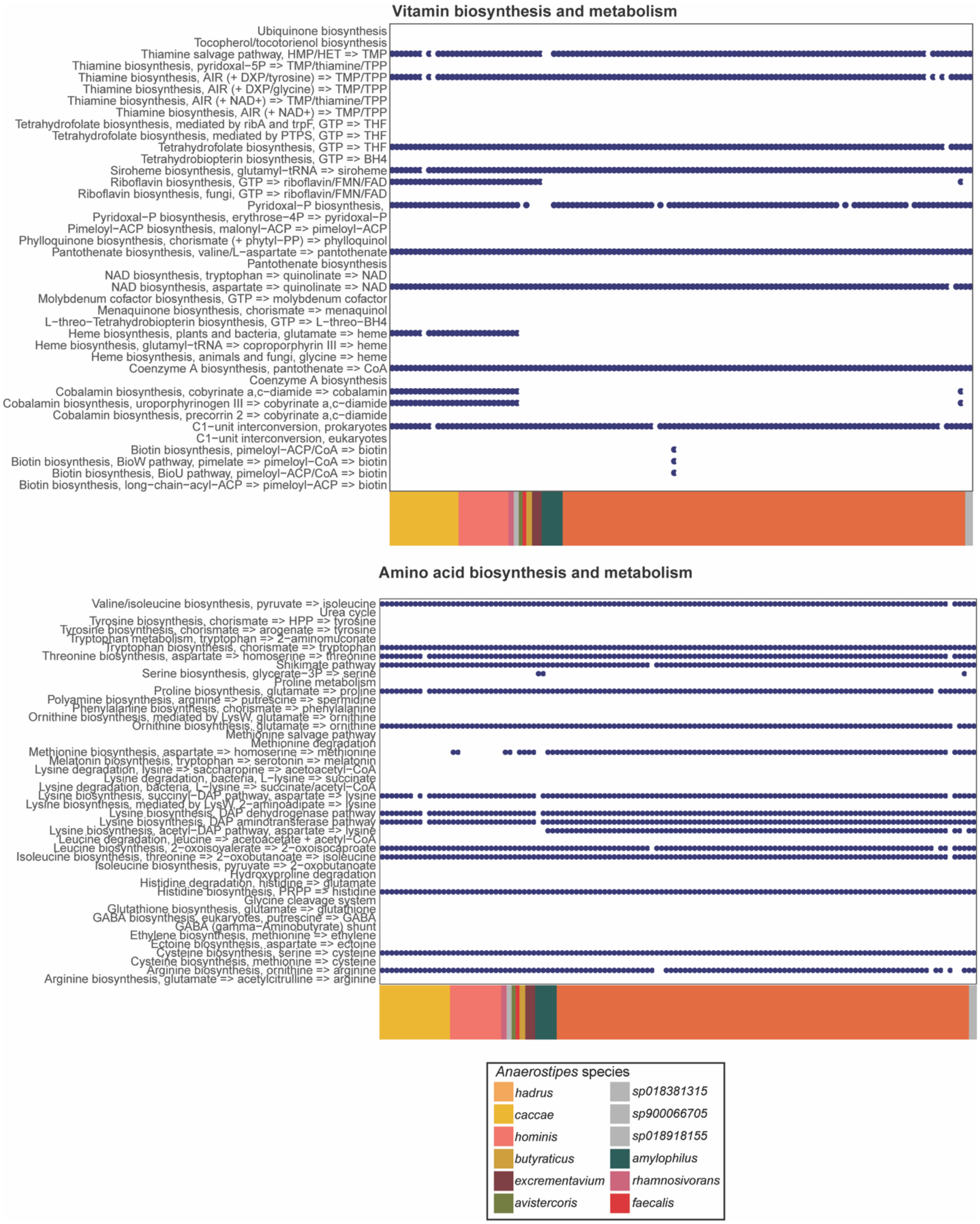


**Figure S4. Module completion for amino acid and vitamin biosynthesis and metabolism pathways identified using Anvi'o.** The color bar in the x axis depict the species in the legend. Presence is depicted by navy blue circle and absence by white.

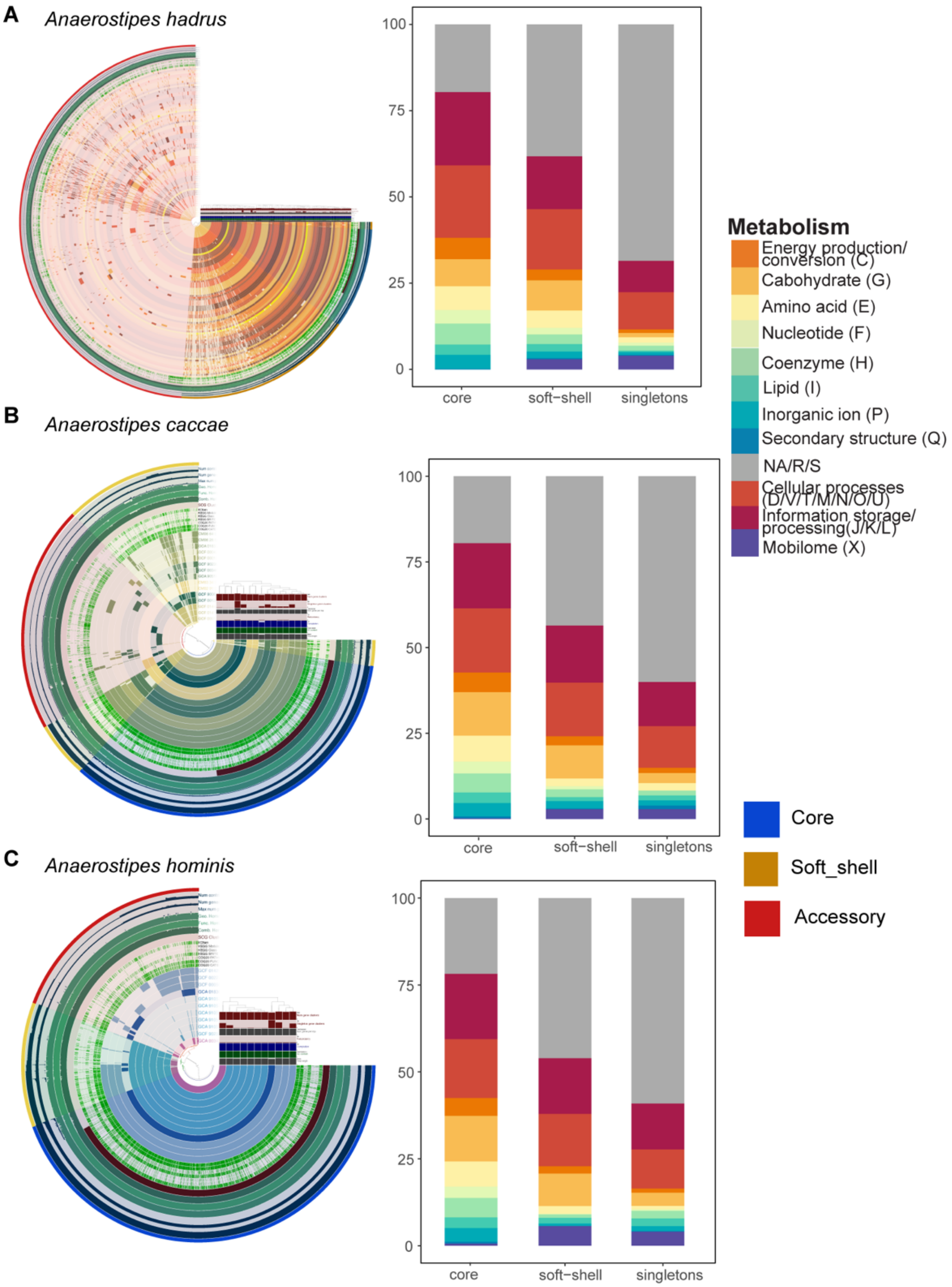


**Figure S5. Pangenomes of *A. hadrus*, *A. caccae* and *A. hominis* with the functional COG categorisation of their genes. A)** *A. hadrus*, **B)** *A. caccae*, and **C)** *A. hominis* pangenomes created using Anvi'o along with relative percentage of the COG categories the genes belong to, in the core (present in all 100% genomes), soft shell (present in 99% of the genomes) and singleton (present only in a single genome) genes.

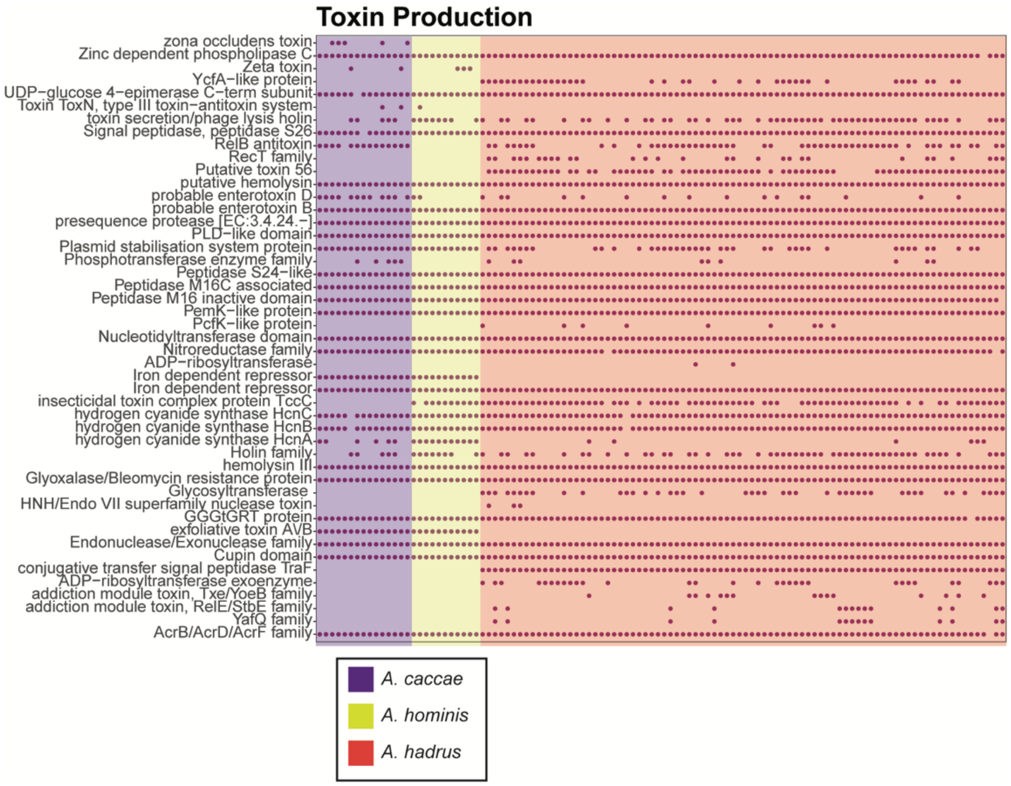


**Figure S6. Virulence in *A. hadrus*, *A. caccae* and *A. hominis*.** Toxins and virulence factors predicted using PathoFact for *A. hadrus*, *A. caccae*, *A. hominis*.
